# A multispecific antibody prevents immune escape and confers pan-SARS-CoV-2 neutralization

**DOI:** 10.1101/2022.07.29.502029

**Authors:** John Misasi, Ronnie R. Wei, Lingshu Wang, Amarendra Pegu, Chih-Jen Wei, Olamide K. Oloniniyi, Tongqing Zhou, Juan I. Moliva, Bingchun Zhao, Misook Choe, Eun Sung Yang, Yi Zhang, Marika Boruszczak, Man Chen, Kwan Leung, Juan Li, Zhi-Yong Yang, Hanne Andersen, Kevin Carlton, Sucheta Godbole, Darcy R. Harris, Amy R. Henry, Vera B. Ivleva, Paula Lei, Cuiping Liu, Lindsay Longobardi, Jonah S. Merriam, Danielle Nase, Adam S. Olia, Laurent Pessaint, Maciel Porto, Wei Shi, Jeremy J. Wolff, Daniel C. Douek, Mehul S. Suthar, Jason Gall, Richard A. Koup, Peter D. Kwong, John R. Mascola, Gary J. Nabel, Nancy J. Sullivan

**Author notes:** Present address: Modex Therapeutics Inc., an OPKO Health Company, Natick, MA 01760, USA. **Equal contribution:** (John Misasi, Ronnie R. Wei, Lingshu Wang, Amarendra Pegu, Chih-Jen Wei). **Equal contribution:** (John R. Mascola, Gary J. Nabel, Nancy J. Sullivan). <u>Corresponding authors</u>: Nancy J Sullivan, Gary J. Nabel.

## Abstract

Despite effective countermeasures, SARS-CoV-2 persists worldwide due to its ability to diversify and evade human immunity^1^. This evasion stems from amino-acid substitutions, particularly in the receptor-binding domain of the spike, that confer resistance to vaccines and antibodies ^2–16^. To constrain viral escape through resistance mutations, we combined antibody variable regions that recognize different receptor binding domain (RBD) sites^17,18^ into multispecific antibodies. Here, we describe multispecific antibodies, including a trispecific that prevented virus escape >3000-fold more potently than the most effective clinical antibody or mixtures of the parental antibodies. Despite being generated before the evolution of Omicron, this trispecific antibody potently neutralized all previous variants of concern and major Omicron variants, including the most recent BA.4/BA.5 strains at nanomolar concentrations. Negative stain electron microscopy revealed that synergistic neutralization was achieved by engaging different epitopes in specific orientations that facilitated inter-spike binding. An optimized trispecific antibody also protected Syrian hamsters against Omicron variants BA.1, BA.2 and BA.5, each of which uses different amino acid substitutions to mediate escape from therapeutic antibodies. Such multispecific antibodies decrease the likelihood of SARS-CoV-2 escape, simplify treatment, and maximize coverage, providing a strategy for universal antibody therapies that could help eliminate pandemic spread for this and other pathogens.

## Main Text

The continued viral replication and transmission of viruses during human pandemics contribute to genetic evolution that can lead to increased pathogenesis and decreased sensitivity to host immunity and antivirals. For SARS-CoV-2, variations in the spike protein have resulted in the appearance of dozens of major variants of concern (VOC). VOCs such as Beta, Delta, Omicron and the newer Omicron sub-lineages contain several to dozens of amino acid variations in their spike protein that have been associated with decreased vaccine and therapeutic antibody efficacy ^17–24^ Among nine antibodies that have previously been, or are currently, approved as COVID-19 therapeutics ^3–16,25^, most have lost potency and/or breadth against evolving and variable circulating variants (**Fig 1a**). The emergence of the original Omicron (BA.1) VOC resulted in high-level resistance to several therapeutic antibodies, with only COV2-2196 (tixagevimab), S309 (sotrovimab) and LY-CoV1404 (bebtelovimab) remaining active (**Fig 1a**) ^17,19,26^. The subsequent BA.2, BA.2.12.1 and BA.4/5 variants showed additional changes in sensitivities: COV2-2196 became inactive against to BA.4/5, REGN10987 regained activity against BA.2 and BA.2.12.1, S309 lost potency against BA.2, BA.2.12.1 and BA.4/5 and COV2-2130 regained potency against BA.2, BA.2.12.1 and BA.4/5 (**Fig 1a**). This “cyclic” variation in potency is a key challenge to maintaining a portfolio of antibody-based therapies against COVID-19.

**Figure 1.**
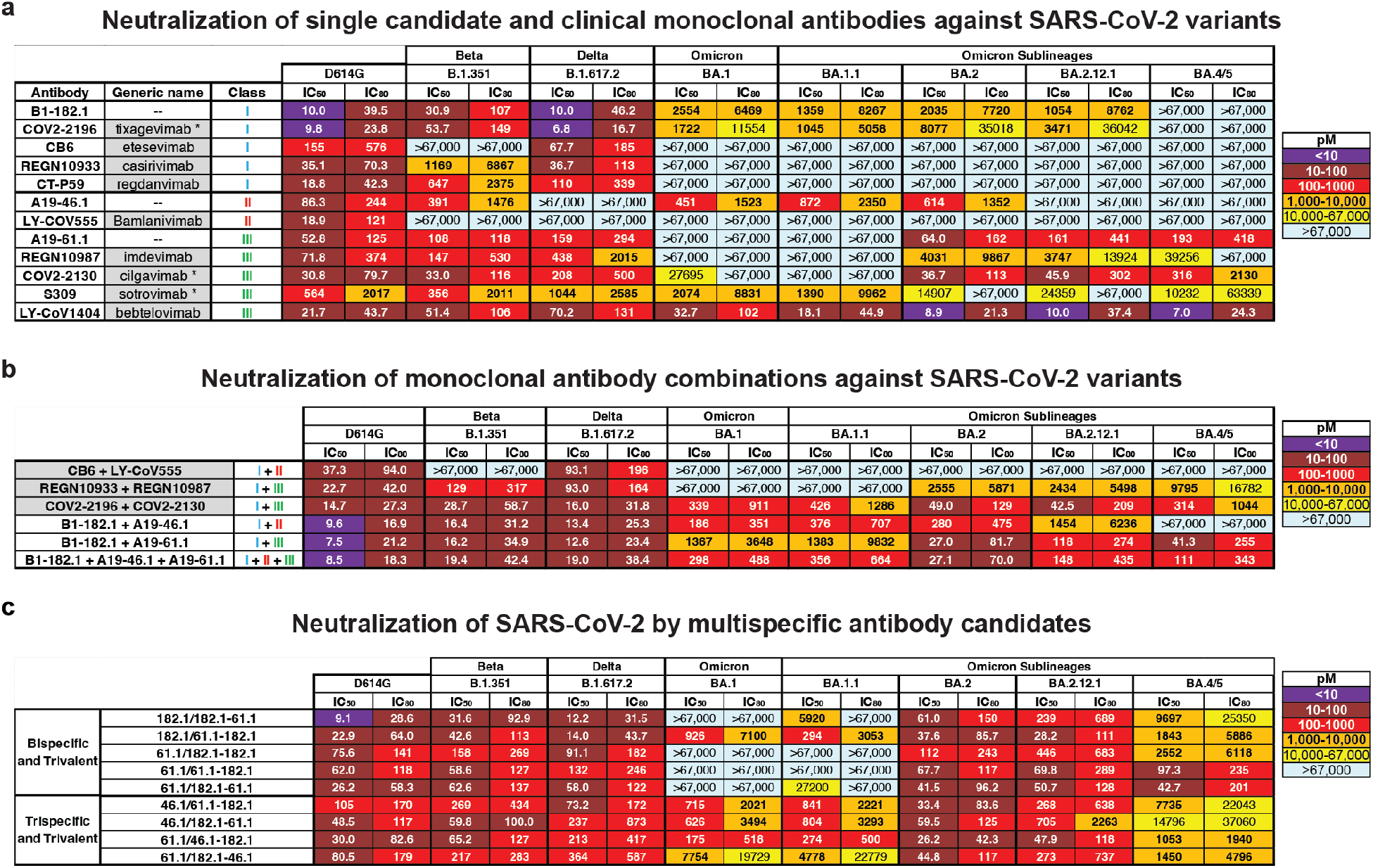
Neutralization of SARS-CoV-2 variants by monoclonal antibodies, antibody cocktails and CODV formatted multispecific antibodies. Neutralization of candidate and expanded access monoclonal antibodies **(a)** and cocktails **(b)** against D614G and the indicated SARS-CoV-2 variants: Beta (B.1.351), Delta (B.1.617.2), Omicron (B.1.1.529 or BA.1) and Omicron sub-lineages. When appropriate, generic names are indicated. Generic names with * indicate the presence of Fc domain mutations in the clinical product that are not found in the experimental versions used in this paper. Class indicates the Barnes RBD classification^13^: class I antibodies bind to the ACE2 binding site when RBD is in the up position; class II bind to the ACE2 binding site when RBD is in the up or down position; class III bind outside the ACE2 binding site when RBD is in the up or down position; and class IV bind outside of the ACE2 binding site when RBD is in the up position. **c,** Neutralization of candidate multispecific antibodies against D614G and the indicated SARS-CoV-2 variants, including Beta (B.1.351), Delta (B.1.617.2), Omicron (B.1.1.529 or BA.1) and Omicron sub-lineages. Neutralization in pM is shown. Ranges are indicated with light blue (>67,000 pM), yellow (>10,000 to ≤67,000 pM), orange (>1,000 to ≤10,000 pM), red (>100 to ≤1,000 pM), maroon (>10 to ≤100 pM), and purple (≤10 pM).

Among the clinical antibodies, only LY-CoV1404 has thus far maintained activity against all prior and current variants. However, as viral evolution continues, resistant variants will develop to any single antibody. Therefore, there is a need to identify antibody therapeutics that can maintain activity in the face of evolving viral antigenic variation. We previously identified three monoclonal antibodies (mAb), B1-182.1, A19-46.1 and A19-61.1 (hereafter termed 182.1, 46.1 and 61.1), that target distinct receptor binding domain (RBD) epitopes and retain high potency and breath against most VOCs ^17,18^: 182.1 displays subnanomolar potency against pre-Omicron VOCs ^17,18^, including Beta and Delta (**Fig 1a**) and maintains potency to BA.1, BA.1.1, BA.2 and BA.2.12.1 that is similar to mAb S309 which showed clinical efficacy against BA.1 ^21,22^; 61.1 has subnanomolar potency against Alpha, Beta and Delta variants but loses activity to Omicron BA.1 and BA.1.1, and then recovers neutralization activity against BA.2, BA.2.12.1 and BA.4/5 ^17,18^ (**Fig 1a**); 46.1 is potently neutralizing against VOC that do not contain substitutions at L452, including Beta, Gamma, BA.1 and BA.1.1 ^17,18^ (**Fig 1a**), but is inactive against BA.2.12.1 and BA.4/5 that have L452Q or L452R. Since these antibodies target distinct epitopes and show a non-overlapping pattern of VOC resistance, it suggested the possibility that a combination of these antibody specificities might provide neutralization breadth against longitudinal SARS-CoV-2 variants.

To test this hypothesis, we combined 182.1 with either 61.1 or 46.1 as two-mAb combinations or with both 61.1 and 46.1 as triple-mAb combination. These were tested for neutralization activity against Beta, Delta, Omicron and Omicron sublineages and results were compared with current clinical antibody cocktails. Consistent with previous reports, the therapeutic cocktails, CB6 + LY-CoV555 and REGN10933 + REGN10987, are unable to neutralize all VOCs (**Fig 1b**)^17,23,24,27^. In contrast, the combination of COV2-2196 + COV2-2130 (e.g., tixagevimab+cilgavimab) maintained subnanomolar potency across all variants albeit with some loss of potency (**Fig 1b**). As we previously showed^17^, the combination of 182.1 + 46.1 provided synergistic neutralization against BA.1 (IC_50_ 186 pM) (**Fig 1b**) compared to the individual components (IC_50_ 2554 and 451 pM) (**Fig 1a**). In addition, we noted similarly potent synergistic neutralization against all Omicron sublineages except for BA.2.12.1 and BA.4/5; for which one or both mAbs lack activity (**Fig 1a, b**). The combination of 182.1 + 61.1 neutralized all variants with IC_50_ values that were equivalent to or better than the parental antibodies (**Fig 1a, b**), and the triple combination of 182.1 + 61.1 + 46.1 neutralized all variants with improved overall potency compared to the double combinations (**Fig 1b**). These results show that combinations of 182.1, 46.1 and 61.1, targeting class I, II and III sites within the SARS-CoV-2 spike RBD, respectively, achieve potent neutralization across all prior and current VOCs.

We previously developed recombinant trispecific antibodies against HIV-1 by combining arms from selected broadly neutralizing antibodies into a single molecule that showed unprecedented potency and breadth ^28^. These antibodies broadly neutralized >98% of circulating virus strains ^28^ and exerted antiviral effects in non-human primates while also mitigating the generation of viral immune escape ^29^. We hypothesized that multispecific antibodies could be designed against SARS-CoV-2 that similarly improve breadth and neutralization reactivity to cover known and evolving antigenic variation. We utilized the cross-over of dual variable (CODV) format comprising one arm heterodimerized with two variable fragment (Fv) domains and the second arm containing a single Fv domain ^30^. We designed nine trivalent anti-SARS-CoV-2 antibodies with bispecific (*i.e*., two antigenic targets) or trispecific (*i.e*., three antigenic targets) reactivity **(Extended Data Figure 1, Extended Data Table 1)**. Five trivalent bispecific antibodies were generated: three containing two Fv182.1 (Fv182) and one Fv61.1 (Fv61) in different positions and two containing two Fv61 and one Fv182 in different positions **(Extended Data Figure 1, Extended Data Table 1)**. Four trivalent trispecific antibodies containing Fv182, Fv61 and Fv46.1 (Fv46) were designed to avoid placing Fv46 and Fv61 in the same CODV arm, since they were previously shown to compete with each other ^18^. To confirm the activity of each Fv component within the trivalent antibodies, we used ELISA to evaluate binding to SARS-CoV-2 RBD proteins containing mutations that specifically eliminate the binding of all but one Fv; specifically, these RBDs contained one or more of the previously identified knockout mutations for Fv182 (F486S), Fv46 (L452R) and Fv61 (K444E) ^18^ **(Extended Data Figure 2a, b)**. We found that each component Fv within the trivalent antibodies recognized its cognate epitope as expected, with equivalent binding to the wildtype RBD where the Fv epitopes were intact (**Extended Data Figure 2c)**. As a further test, we assessed the ability of each antibody to neutralize pseudoviruses with spike protein point mutations that eliminated binding of a single Fv component and showed that each trivalent multispecific mAb maintained neutralization via the remaining Fvs (**Extended Data Figure 2d**). Taken together, these findings verified that the Fvs within each multispecific antibody were functioning as expected.

We next evaluated the ability of the trivalent antibodies to neutralize the SARS-CoV-2 ancestral D614G, and the Beta, Delta, BA.1, BA.1.1, BA.2, BA.2.12.1 and BA.4/5 variants. For D614G, Beta and Delta variants, all of the multispecifics neutralized with subnanomolar IC_50_s (**Fig 1c**). Among the trivalent dual recognition (bispecific) antibodies that included 182.1 and 61.1 in different configurations, we observed that the antibody with the 182.1/61.1-182.1 configuration maintained subnanomolar affinity against BA.1 and BA.1.1 (**Fig 1c**). These data indicate that for BA.1 and BA.1.1, the positioning of the Fv domain within a multispecific can impact neutralization potency. All of these bispecific antibodies displayed subnanomolar neutralization against BA.2 and BA.2.12.1 likely because, among the Fvs, the parental 61.1 antibody regains potent neutralization against these lineages. Interestingly, we found that having two copies of Fv61, regardless of position, significantly improved BA.4/5 neutralization >100-fold over other bispecifics antibodies containing a single Fv61 **(Fig 1c)**. For the trispecific antibodies, all variants tested were neutralized with pico- to nanomolar potency. Against D614G, Beta and Delta variants IC_50_s ranged between 30 and 364 pM **(Fig 1c)**. For BA.1, BA.1.1 and BA.4/5, neutralization by 61.1/46.1-182.1 occurred with IC_50_s of 175 pM, 274 pM and 1053 pM **(Fig 1c)**, respectively, and notably for BA.1, with higher potency than the clinically effective antibody sotrovimab (IC_50_ 2074 pM) **(Fig 1a).** The 1-2 log higher potency of 61.1/46.1-182.1 for BA.1 and BA.1.1 compared to 61.1/182.1-46.1 suggests that the relative positions of 182.1 and 46.1 in the CODV arm are impacting on trispecific mAb potency. In contrast for BA.2 and BA.2.12.1, all four trispecific antibodies neutralized with IC_50_ between 26 and 705 pM, with 61.1/46.1-182.1 again having the highest potency **(Fig 1c)**. These data indicate that a multispecific antibody with a precise configuration of Fv domains broadly neutralizes diverse strains (including strains that did not exist when the mAbs were made), suggesting that inter-epitope engagement increases antibody potency and breadth.

We note that the most potent and broad antibody, trispecific 61.1/46.1-182.1, contains the same component Fvs in its CODV arm as trispecific 61.1/182.1-46.1, yet neutralizes BA.1 and BA.1.1 15 to 40-fold better than 61.1/182.1-46.1 **(Fig 1c)**. To better understand how differences in the positions of Fv46 and Fv182 in the CODV arm of these molecules led to differences in potency, we used negative stain-electron microscopy (NSEM) to compare the binding of purified CODV 46.1-182.1 or 182.1-46.1 to D614G spike trimer proteins containing mutations that eliminate binding to either Fv46 or Fv182 **(Fig. 2a)**. Consistent with the 182 mAb epitope being at the distal tip of RBDs when they are in the up position, and thus more accessible for binding by Fv, the relative position of Fv182 within the CODV arm did not influence its ability to bind to the trimer **(Fig 2b, 2c)**. However, for Fv46 we noted CODV position-dependent binding to spike trimers. In particular, when Fv46 is in the outer position **(Fig 2d, leftmost panels)**, no NSEM binding classes were observed against K444E/F486S spike protein that should be recognized by Fv46 but is unable to bind Fv182 (**Fig 2d, rightmost panels)**. Since CODV Fv46 binding to soluble RBD was not impacted **(Extended Data Figure 2c)**, these results suggests that the outer position of the CODV is not compatible with binding of Fv46 to RBD domains contained within trimeric spike proteins. On the other hand, when Fv46 is in the inner position **(Fig 2e, leftmost panels)**, NSEM class averages show that Fv46 binds K444E/F486S spike protein (**Fig 2e, center-right)**. Representative models of the binding mode observed in the NSEM micrographs show that when Fv46 is in the outer position (**Fig 2e, right panel)**, its angle of approach allows the CHCL1-Fv182 portions of the CODV to be positioned away from the spike trimer, thereby avoiding potential clashes with the spike trimer. Since NSEM of CODV 46.1-182.1 revealed that both Fv domains can bind spike trimers, we hypothesized that the CODV 46.1-182.1 alone would be sufficient to cross-link Omicron spike trimers. Indeed, NSEM micrographs revealed that aggregates were formed when CODV 46.1-182.1 was incubated with Omicron spike trimers **(Fig 2e)**. Taken together, these results illustrate how in the context of spike trimers, both the location of epitopes within RBD and CODV position can impact binding, aggregative potential and neutralization. Specifically, Fvs with an angle of approach vertical to the trimer apex such as Fv182 are likely to have greater flexibility for positioning with trivalent mAb designs, likely due to freedom from steric constraints, and consistent with higher potency neutralization by this Fv. In contrast, Fv46 has a lateral or angled approach suggesting that antibodies that bind with this angle of approach may be subject to steric constraints that are revealed by position-dependence of the Fv for optimal engagement and neutralization.

**Figure 2.**
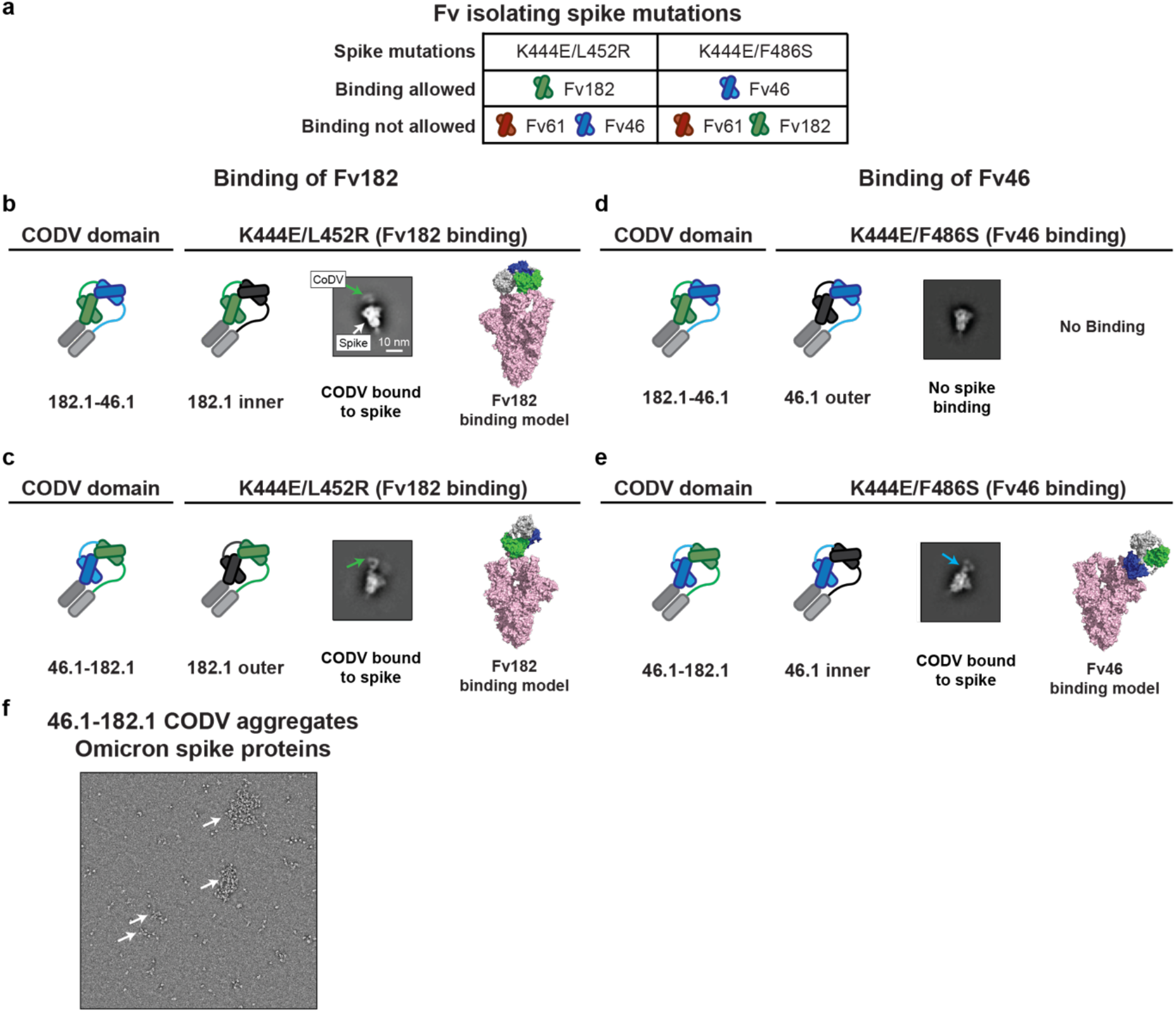
Identification of potential binding modes of CODV to SARS-CoV-2 spike by negative stain-electron microscopy. **a,** Combination of RBD mutations on the SARS-CoV-2 spike designed to distinguish different binding modes of CODV. Mutations were made in spike in the spike trimer. The domains are colored green, blue, red and gray for Fv182.1, Fv46.1, Fv61.1 and constant domain, respectively. **b-e,** Evaluation of CODV Fv182 (**b** and **c**) binding to K444E/L452R or CODV Fv46 (**d** and **e**) binding to K444E/F486S. Shown at left in each panel is a schematic of the CODV domain being evaluated in the panel. The center-left subpanel indicates the Fv and position being evaluated (i.e., inner or outer). For clarity, the Fv domain that is not able to bind to the mutant is colored black in the subpanels. The center-right subpanel shows the negative stain electron micrograph class averages. Fv182 are indicated with a green arrow and Fv46 with a blue arrow. The white scale bar represents 10 nm in panel **c** and applies to each micrograph. The rightmost subpanel is a representative model of the binding mode observed in the micrograph. Panels **b** and **c** show that irrespective of position, both 182.1-46.1 and 46.1-182.1 are able to bind to spike protein that only allows binding by the Fv182. Panel **d**, shows that 182.1-46.1 (46.1 outer) is unable to bind spike protein that only allows binding by the Fv46.1. Panel **e**, shows that 46.1-182.1 (46.1 inner) is able to bind spike that only allows binding by Fv46. **f,** CODV 46.1-182.1 induced aggregation of the Omicron spike. Large clusters of aggregation were observed in the negative stain EM field, suggesting the CODV 46.1182.1 can efficiently promote inter-spike crosslinking.

The perpetuation of the COVID-19 pandemic due to waves of infection by emerging VOCs has demonstrated the impact of viral evolution and in particular, the impact of antigenic changes in the spike protein, on virus persistence. We previously showed that replication-competent vesicular stomatitis virus (rcVSV) pseudotyped with the SARS-CoV-2 spike can rapidly mutate to escape neutralization by the individual antibodies 182.1, 46.1 or 61.1 ^18^. We therefore sought to compare the potential for escape from the trispecific antibody with the broadest and most potent neutralization activity, 61.1/46.1-182.1, against the triple antibody cocktail or the individual mAbs, 182.1 and LY-CoV1404. Under conditions where 182.1 and LY-CoV1404 fully escaped antibody neutralization (i.e., >20% cytopathic effect at 333,333 pM) within 2-3 rounds of repeated infection *in vitro* **(Fig 3)**, we found that an equal mixture of 182.1 + 61.1 + 46.1 slowed acquisition of the escaped phenotype, but gradually lost neutralization potency during 7 rounds of infection (**Fig 3**). For trispecific, 61.1/46.1-182.1, there was no observed escape in two independent experiments; though from round 1 to round 2 of replication there was a modest shift in the antibody concentration (107 pM) required to maintain <20% CPE, in line with neutralization potency determinations that would not be expected to fully suppress viral growth **(Fig 1c and 3)**. The ability of this trispecific Ab to mitigate neutralization escape corresponds with the observation that the CODV arm of 61.1/46.1-182.1 can also cross-link and aggregate spikes due to both Fv components being able to bind RBD (**Fig 2c, 2e and 2f)**. Since the individual mAbs and mAb cocktail would not be expected to crosslink spikes, these results suggest that the potency and improved escape mitigation of 61.1/46.1-182.1 over the antibody cocktail is mediated by the ability to the trispecific Ab to engage and aggregate multiple trimers.

**Figure 3.**
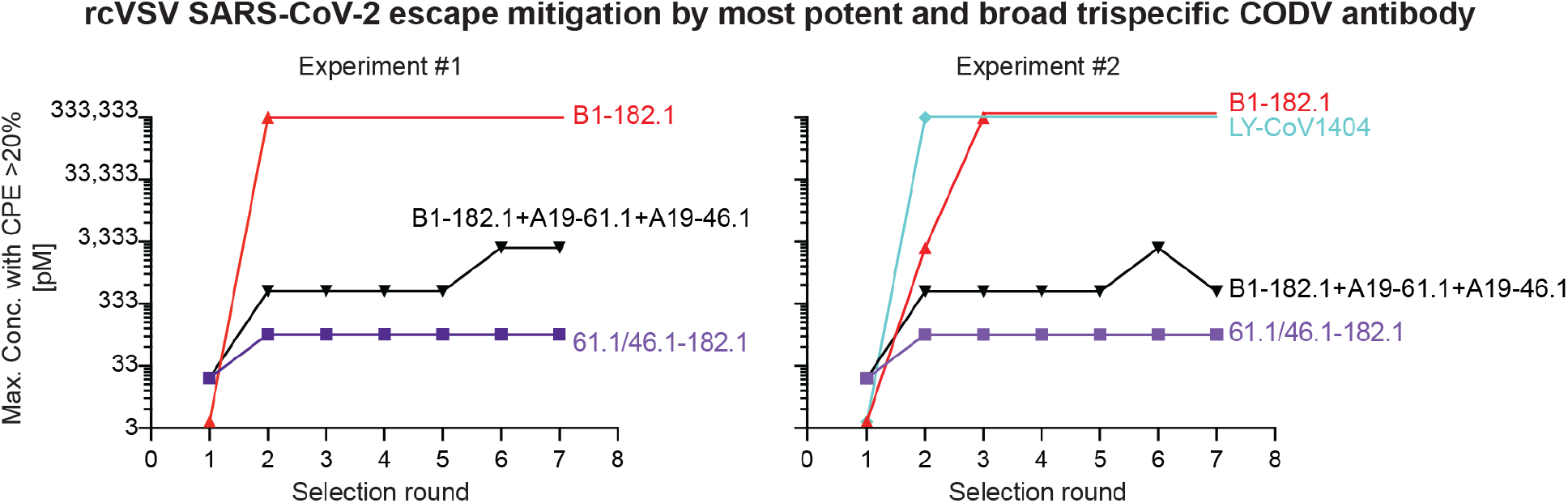
Mitigation of rcVSV-SARS-CoV-2 escape by 61.1/46.1-182.1 and an antibody cocktail. Replication competent VSV (rcVSV) bearing SARS-CoV-2 WA-1 spike protein was incubated with the indicated antibodies at 5-fold increasing concentrations (34 × 10^-3^ to 333,333 pM) and added to Vero cells. The appearance of viral growth, as indicated by the presence of >20% CPE, was determined after 3 days and supernatant from the highest antibody concentration with viral growth was passaged forward into a new selection round under the same conditions. Once viral growth appeared at 333,333 pM, the antibody was considered fully escaped and supernatant was no longer passaged forward. Data is graphed as the highest concentration of antibody at which viral growth was noted in each selection round. Each graph represents an independent experiment.

We next designed a tetravalent trispecific antibody, 46.1-182.1v/61.1-182.1v, with single Fv46 and Fv61 domains and two Fv182v domains; Fv182v is a stabilized variant of Fv182. We found that 46.1-182.1v/61.1-182.1v showed similar breadth and potency to 61.1/46.1-182.1 **(Extended Data Figure 3)**. While therapeutic COVID-19 antibodies can be authorized for use against Omicron variants based on clinical pharmacokinetic data, along with the non-clinical viral neutralization data for Omicron variants ^31^, we nonetheless sought to determine the capacity of 46.1-182.1v/61.1-182.1v to protect Syrian golden hamsters from infection by BA.1, BA.2 and BA.5 **(Figure 4a)**. Consistent with previous reports of attenuated BA.1 pathogenesis in hamsters^32^, we found that PBS treated animals showed a delay in weight loss (day 5) (**Fig 4b)**. For the 46.1-182.1v/61.1-182.1v treated animals, we noted that the hamsters continued to gain weight throughout the study **(Fig 4b)**. We next assessed viral loads in the lung tissue of PBS and antibody treated animals and found that while PBS treated animals show >7-logs of virus per gram (TCID_50_/g) up until day 4, antibody treated animal had very low to undetectable virus in the assay **(Fig 4c).** Next, we infected BA.2 and BA.5 hamsters infected with sequence-verified BA.2 (2×10^4^ PFU) or BA.5 SARS-CoV-2 (1×10^5^ PFU). In contrast to a recent report that noted no significant weight loss in hamsters infected with BA.2 and BA.5^33^, we observed 7-10% weight loss beginning on day 3 in infected, untreated animals. In contrast, antibody-treated animals continued to gain weight throughout the study **(Fig 4d, f)**. Similar to BA.1, the viral titer for BA.2 was >7-logs of virus per gram on days 2 and 4 in PBS treated animals and no virus growth was observed in the lungs of treated animals **(Fig 4 e)**. For BA.5, viral titer exceeded 8-logs on day 2 and antibody treated animals showed a significant 13-fold lower titer on day 4 relative to PBS treated animals **(Fig 4 g)**. Together these data suggest that 46.1-182.1v/61.1-182.1v has breadth, potency and *in vivo* efficacy against Omicron and Omicron sublineages with distinct amino acid substitutions that mediate escape therapeutic antibodies.

**Figure 4.**
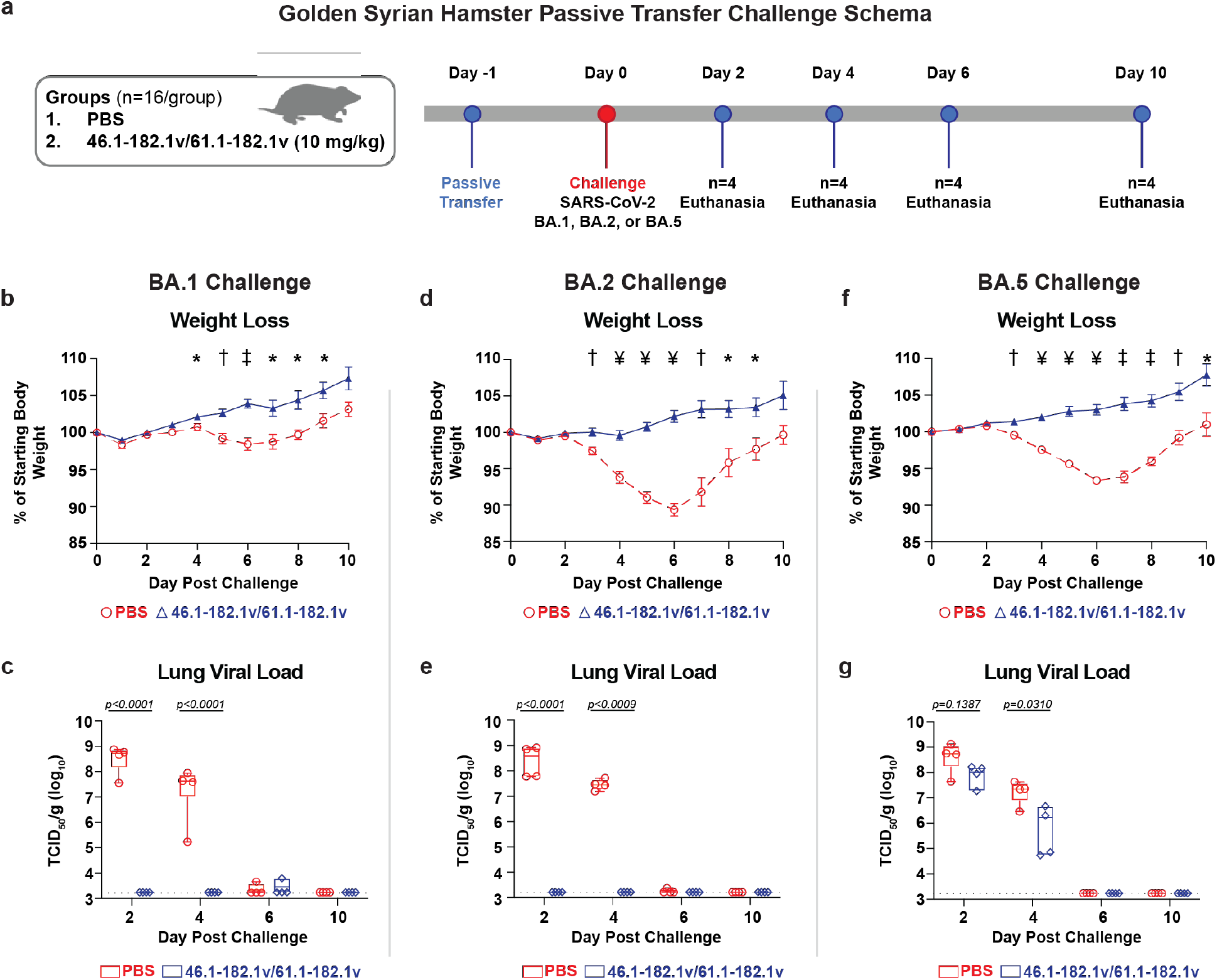
A trispecific antibody protects Syrian hamsters from SARS-CoV-2 variant BA.2. **a**, 46.1-182.1v/61.1-182.1v was passively transferred to Syrian hamsters intraperitoneally 24 hours prior to challenge with either 1×10^5^ PFU of BA.1, 2×10^4^ PFU of BA.2 or 1×10^5^ PFU of BA.5 SARS-CoV-2 Omicron variants of concern. For each virus, an additional group of hamsters received an equal volume of PBS in the same manner and served as controls. All animals were weighed daily to monitor weight loss and 4 animals per group were euthanized at days 2, 4, 6, and 10 to assess viral loads in the lung. **(b, d, f)** Hamsters were monitored daily for weight loss over 10 days. Symbols and error bars represent means and SEM, respectively. An unpaired, two-tailed student *t*-test was used to determine significance at each timepoint between treated and untreated animals. Significant *p*-values are indicated as: * (≤0.05), ^†^ (≤0.01), ^‡^ (≤0.001) and ^¥^ (≤0.0001). (**c, e, g**) Viral load in the lung of BA.1 (c), BA.2 (e) and BA.5 (g) challenged hamsters were quantified by TCID_50_ per gram. Each symbol represents an individual animal and may overlap for equivalent values. Boxes and horizontal bars denote the IQR and medians, respectively; whisker end points are equal to the maximum and minimum values. Dotted lines indicate the assay’s lower limit of detection. An unpaired, two-tailed student t-test was used to determine significance at each timepoint between treated and untreated animals. Significant p-values are indicated on graph.

In this manuscript, we developed multispecific antibodies that target independent epitopes on the viral spike. These antibodies are highly potent and maintain breadth against VOCs with evolving patterns of antigenic variation, including the most recent Omicron sublineages. The combination of three antibody specificities in a precise orientation within one molecule ensured that the antibody could neutralize VOC with IC_50_s in a range similar to, or better than, clinically active antibodies, even when only one antibody Fv domain was active. In addition, the trispecific antibody mitigated virus escape *in vitro* under conditions where highly potent and broad single monoclonal antibodies could not, even when combined as cocktails. While the antibodies in this study were initially isolated prior to the emergence of Omicron, the strategy of rationally choosing antibody specificities that target distinct antigenic sites on spike, and that are differentially impacted by VOC mutations, allowed the generation of broadly active molecules that neutralized and had protective efficacy in hamsters against variants that were unforeseen at the time of their design. Furthermore, the inclusion of multiple specificities on the same molecule has the potential of additive or synergistic binding and restriction of pathways for viral escape. Finally, the results in this report suggest the possibility that vaccine antigens targeting the functionally constrained epitopes contained in these trispecific antibodies might increase breadth and potency against current and future variants using a single protein with simplified clinical development.

## Methods

### Design of trispecific antibodies

The trispecific antibody format follows the previously described design configuration ^28^. Briefly, the bispecific antibody arm ^30^ is heterodimerized with a conventional antibody arm, or alternatively, two bispecific antibody arms are heterodimerized using knob-in-hole ^34^ mutations in the CH3 domain of IgG1 Fc. Specifically, three classes of anti-SARS-CoV-2 neutralizing antibodies targeting RBDs B1-182.1 (class I), A19-46.1 (class II), and A19-61.1 (class III) were tested for possible combinations including each antibody Fv orientation and copy number to achieve bi- or tri-specificity. These combinations were tested for best neutralizing breadth/potency. In addition, “Knob” (S354C/T366W), “Hole” (Y349C /T366S/L368A/Y407V) ^34^, and “LS” (M428L/N434S) ^35^ mutations were engineered into CH3 of the monospecific, bi- or trispecific Fc region.

### Synthesis, cloning and expression of multispecific antibodies

After design of the amino acid sequences for each multispecific antibody, the four genes for each multispecific antibody were synthesized using human preferred codons (GenScript) and cloned into eukaryotic expression vectors. For each multispecific antibody expression, equal amounts of the 4 plasmid DNAs were transfected into Expi293 cells (ThermoFisher) using Expi293 transfection reagent (ThermoFisher) as previously reported ^28^. The transfected cells were cultured in shaker incubator at 120 rpm, 37 °C, 9% CO2 for 4~5 days. Culture supernatants were harvested and filtered, the multispecific antibodies were purified over a Protein A (GE Health Science) column. Each multispecific antibody was eluted with IgG elution buffer (Pierce), immediately buffer exchanged with PBS and concentrated using Centricon Plus-70 (Millipore Sigma) membrane filter unit. After concentration, each multispecific antibody was applied to a Superdex 200 16/600 size exclusion column (Cytiva) to remove aggregates and different species in the preparation. The fractions were then analyzed on reduced and non-reduced SDS-PAGE to identify the fractions that contained the monomeric multispecific antibody before combining them. The pooled fractions were then further concentrated, aliquoted and analyzed by SDS-PAGE as well as an analytical SEC column (Superdex 200 16/600) to verify purity. Molecular weight, extinction coefficient and predicted pI were determined using Geneious Prime (Biomatters Ltd.)

To make the CODV from multispecific IgG, the FabALACTICA protease (Genovis) was used to digest the IgG for 16 hrs at room temperature. The digestion mixture was then incubated with protein A resin to remove Fc and undigested IgG, the flowthrough and PBS wash of the protein A column that contained the Fab and CODV fragments was collected, concentrated and further purified with size exclusion column (Superose 6 10/300, Cytiva).

The plasmids encoding 46.1-182.1v/61.1-182.1 were transfected into Expi293 cells using ExpiFectamine 293 (ThermoFisher) following the manufacturer’s protocol. The transfected cells were then cultured in shaker incubator at 120 rpm, 37 °C, 9% CO2 for 4 days and transferred to a second shaker incubator at 32°C one day before harvest. The 46.1-182.1v/61.1-182.1 antibody was purified by using Protein A Sepharose (Cytiva), followed by size-exclusion chromatography using a Superdex 200 16/600 column (Cytiva). Purified 46.1-182.1v/61.1-182.1 was stored in a buffer containing PBS, pH 6.0 and 5% sucrose.

### Full-length S constructs

Codon optimized cDNAs encoding full-length S from SARS-CoV-2 (GenBank ID: QHD43416.1) were synthesized, cloned into the mammalian expression vector VRC8400 ^36,37^ and confirmed by sequencing. S containing D614G amino acid change was generated using the wt S sequence. Other variants were made by mutagenesis using QuickChange lightning Multi Site-Directed Mutagenesis Kit (cat # 210515, Agilent) or via synthesis and cloning (Genscript) as previously reported ^18,38^. The S variants tested are B.1.351 (L18F, D80A, D215G, (L242-244)del, R246I, K417N, E484K, N501Y, A701V), B.1.1.7 (H69del, V70del, Y144del, N501Y, A570D,D614G, P681H, T716I, S982A, D1118H), B.1.617.2 (T19R, G142D, E156del, F157del, R158G, L452R,T478K, D614G, P681R, D950N), B.1.1.529 or BA.1 (A67V,H69del, V70del, T95I, G142D, V143del, Y144del,Y145del, N211del, L212I, ins214EPE, G339D,S371L, S373P, S375F, K417N, N440K, G446S,S477N, T478K, E484A, Q493R, G496S, Q498R,N501Y, Y505H, T547K, D614G, H655Y, N679K,P681H, N764K, D796Y, N856K, Q954H, N969K,L981F), BA.1.1 (A67V,H69del, V70del, T95I, G142D, V143del, Y144del,Y145del, N211del, L212I, ins214EPE, G339D, R346K, S371L, S373P, S375F, K417N, N440K, G446S,S477N, T478K, E484A, Q493R, G496S, Q498R,N501Y, Y505H, T547K, D614G, H655Y, N679K,P681H, N764K, D796Y, N856K, Q954H, N969K,L981F), BA.2 (T19I, L24-, P25-, P26-, A27S, G142D, V213G, G339D, S371F, S373P, S375F, T376A, D405N, R408S, K417N, N440K, S477N, T478K, E484A, Q493R, Q498R, N501Y, Y505H, D614G, H655Y, N679K, P681H, N764K, D796Y, Q954H, N969K), BA.2.12.1 (T19I, L24-, P25-, P26-, A27S, G142D, V213G, G339D, S371F, S373P, S375F, T376A, D405N, R408S, K417N, N440K, L452Q, S477N, T478K, E484A, Q493R, Q498R, N501Y, Y505H, D614G, H655Y, N679K, P681H, S704L, N764K, D796Y, Q954H, N969K) and BA.4/5 (T19I, L24-, P25-, P26-, A27S, H69-, V70-, G142D, V213G, G339D, S371F, S373P, S375F, T376A, D405N, R408S, K417N, N440K, L452R, S477N, T478K, E484A, F486V, Q498R, N501Y, Y505H, D614G, H655Y, N679K, P681H, N764K, D796Y, Q954H, N969K). These full-length S plasmids were used for pseudovirus neutralization assays.

### Pseudovirus neutralization assay

S-containing lentiviral pseudovirions were produced by co-transfection of packaging plasmid pCMVdR8.2, transducing plasmid pHR’ CMV-Luc, a TMPRSS2 plasmid and S plasmids from SARS-CoV-2 variants into 293T cells using Lipofectamine 3000 transfection reagent (L3000-001, ThermoFisher Scientific, Asheville, NC) ^39,40^. 293T-ACE2 cells (provided by Dr. Michael Farzan) were plated into 96-well white/black Isoplates (PerkinElmer, Waltham, MA) at 7,500 cells per well the day before infection of SARS CoV-2 pseudovirus. Serial dilutions of mAbs were mixed with titrated pseudovirus, incubated for 45 minutes at 37°C and added to cells in triplicate. Following 2 h of incubation, wells were replenished with 150 ml of fresh media. Cells were lysed 72 h later, and luciferase activity was measured with MicroBeta (Perking Elmer). Percent neutralization and neutralization IC_50_s, IC80s were calculated using GraphPad Prism 8.0.2.

### Antibody binding to RBD mutation proteins by ELISA

MaxiSorp Immuno plates (Thermo Fisher) plates were coated with 1 μg/ml of SARS-CoV-2 WA-1 RBD or RBD with single, double or triples mutations (F486S, E444K, L452R, F486S/E444K, F486S/L452R and F486S/E444K/L452R) in PBS at 4 °C overnight. After standard washes and blocking, plates were incubated with serial dilutions of antibody for one hour at room temperature. Anti-human IgG Fc g-specific horseradish peroxidase conjugates (Jackson Laboratory) was used to detect binding of antibody to the RBD proteins. The plates were then washed and developed with 3,5,3’5’-tetramethylbenzidine (TMB) (KPL, Gaithersburg, MD). After stopped with 1N H2SO4 (Fisher), OD 450 nM was read with a SpectraMax Plus microplate reader (Molecular Device).

### rcVSV SARS-CoV-2 antibody escape assay

Selection of virus escape variants was conducted as previously described ^18^. Briefly, an equal volume of clonal population of replication competent vesicular stomatitis virus (rcVSV) with its native glycoprotein replaced by the Wuhan-1 spike protein (rcVSV SARS-CoV-2) ^41^ at an MOI of 0.01 was mixed with serial dilutions of antibodies (5-fold) in cell media to give the desired final antibody concentration. Antibody cocktails were mixed at equal ratios. Virus:antibody mixtures were incubated at 37°C for 1 hour prior to being added to Vero E6 cells. Virus replication was assessed 72hrs after infection in the presence of selected antibodies. Supernatants from the well with the highest concentration of antibody which showed evidence of viral replication (>20% cytopathic effect) was passaged into the subsequent rounds of selection. Infection, monitoring, and collection of supernatants was performed as in the initial round.

### Expression and Purification of Soluble Spike Constructs

The soluble S protein mutants were made in a background of the HexaPro stabilization of the spike ^42^, incorporating D614G/K444E/L452R and D614G/K444E/F486S, and the protein was produced as previously described ^43^. One liter of Freestyle cells was transfected with 1mg of SARS-CoV-2 spike DNA premixed with 3mL of Turbo293 Transfection Reagent. The cells were grown for 6 days at 37°C, after which the supernatant was collected by centrifugation and filtration. The supernatant was incubated with nickel resin for 1 hour at room temperature, and then the resin was washed with 1X PBS pH 7.4. The spike was eluted with 20mM HEPES pH 7.5, 200mM NaCl, 300mM imidazole and concentrated before loading onto a Superdex S-200 gel filtration column equilibrated in 1X PBS pH 7.4. The trimer containing peak was collected, concentrated to 1mg/ml, flash frozen in liquid nitrogen, and stored at − 80°C until use.

### Negative Stain Electron Microscopy

SARS-CoV-2 spike proteins were mixed with CODV fragments at a molar ratio of 1:1.2 and incubated at room temperature for 10 min and then diluted to a concentration of approximately 0.02 mg spike/ml with 10 mM HEPES, pH 7.4, 150 mM NaCl. To make a grid, 4.8-μl of the diluted sample was placed on a freshly glow-discharged carbon-coated copper grid for 15 s. The drop on grid was then wicked away with filter paper, and the grid was washed and wicked three times. Same volume of 0.75% uranyl formate was added to the grid to negatively stain protein molecules adsorbed to the carbon and immediately wicked away. After three times staining, the grid was allowed to air-dry. Datasets were collected using a Thermo Scientific Talos F200C transmission electron microscope equipped with a Ceta camera at 200 kV. The nominal magnification was 57,000x, corresponding to a pixel size of 2.53 Å, and the defocus was set at −2 μm. Data was collected automatically using EPU. Single particle analysis was performed using CryoSPARC 3.0.

### Syrian Hamster Model, SARS-CoV-2 Infection and Passive Transfer

All experiments were conducted according to NIH regulations and standards on the humane care and use of laboratory animals as well as the Animal Care and Use Committees of Bioqual, Inc. (Rockville, Maryland). Six- to eight-week-old male Syrian hamsters (Envigo) were housed at Bioqual, Inc. Multispecific antibody was diluted into PBS to achieve a dose of 10 mg/kg in a 1-2 mL volume based on the weights and was passively transferred to the hamsters by injection into the peritoneal cavity 24 hours prior to challenge. Hamsters were infected with SARS-CoV-2 variants by injecting 100 μl of virus diluted in PBS into the nares and split between both nostrils. Weight changes and clinical observations were collected daily for all studies. Four hamster per group were euthanized at days 2, 4, 6 and 10 to collect the lung for quantification of viral load. The BA.1 SARS-CoV-2 variant used in this study was previously described^44^. The BA.2 SARS-CoV-2 variant was obtained from the Biodefense and Emerging Infections Research Resources Repository (BEI, NR-56522). The BA.5 SARS-CoV-2 variant was propagated by the laboratory of Mehul S. Suthar in Vero-TMPRSS2 cells and sequence confirmed prior to challenge studies. All SARS-CoV-2 stocks were titrated in hamsters to confirm pathogenicity prior to use in our studies.

### TCID_50_ quantification of SARS-CoV-2

Tissues were weighed, placed into pre-labeled Sarstedt cryovials, and snap-frozen until needed. Prior to testing, the tissues were thawed and homogenized using a hand-held tissue homogenizer. The samples were spun down to remove debris and supernatants were assayed. TCID_50_ was quantified as previously described^45^. Briefly, Vero-TMPRSS2 cells were incubated at 37 °C, 5% CO_2_ overnight. The following day the medium was aspirated and replaced with fresh medium. The samples were serially diluted ten-fold for quantification. Positive (virus stock of known infectious titer in the assay) and negative (medium only) control wells were included in every assay. The plates were incubated at 37 °C, 5.0% CO_2_ for 4 days. The cell monolayers were visually inspected for cytopathic effect. TCID_50_ values were calculated using the Reed–Muench formula. TCID_50_ values were log transformed prior to statistical analysis.

### Data and Code availability

All data is available in the main text or the supplementary materials. Original materials in this manuscript are available under a materials transfer agreement with the National Institutes of Health.

### Statistical analysis

Comparisons between groups are based on two-sided unpaired t-tests. Viral loads are log-transformed and reported as box-and-whisker plots with the line depicting the median and the box extending from the 25th to 75th percentiles. All analyses were conducted using GraphPad Prism version 9.0.

## Acknowledgements

We wish to acknowledge Barney Graham, for insightful discussions, and Yilie Li and Melissa Resto for insightful discussions on antibody quality control assays. This work was funded by the Intramural Research Program of the Vaccine Research Center, NIAID, NIH and by ModeX Therapeutics, Inc

## Author Contributions

J.M., L.W., A.P., R.R.W., C-J.W., Z-y.Y., J.I.M, T.Z. and J.L. designed experiments and analyzed data. J.M., A.P., L.W., T.Z., J.I.M, M.C., B.Z. O.K.O., M.B., Y.Z., E.S.Y., M.C., K.L., J.J.W., V.B.I., A.R.H, S.G., P.L., R.W., C.J.W., J.S.M., H.A., D.N., L.P., M.P., Z-y.Y. and J.L. performed experiments. W.S., A.S.O., D.R.H., M.C. and C.L. produced proteins, antibodies and other reagents. J.M., K.C., R.A.K., D.D., M.S., J.G., J.R.M, P.D.K., G.J.N. and N.J.S supervised experiments. J.M., P.D.K., J.R.M., G.J.N. and N.J.S. wrote the manuscript with help from all authors.

