## Extended Data for "A multispecific antibody prevents immune escape and confers pan-SARS-CoV-2 neutralization"

### Extended data figure/table legends

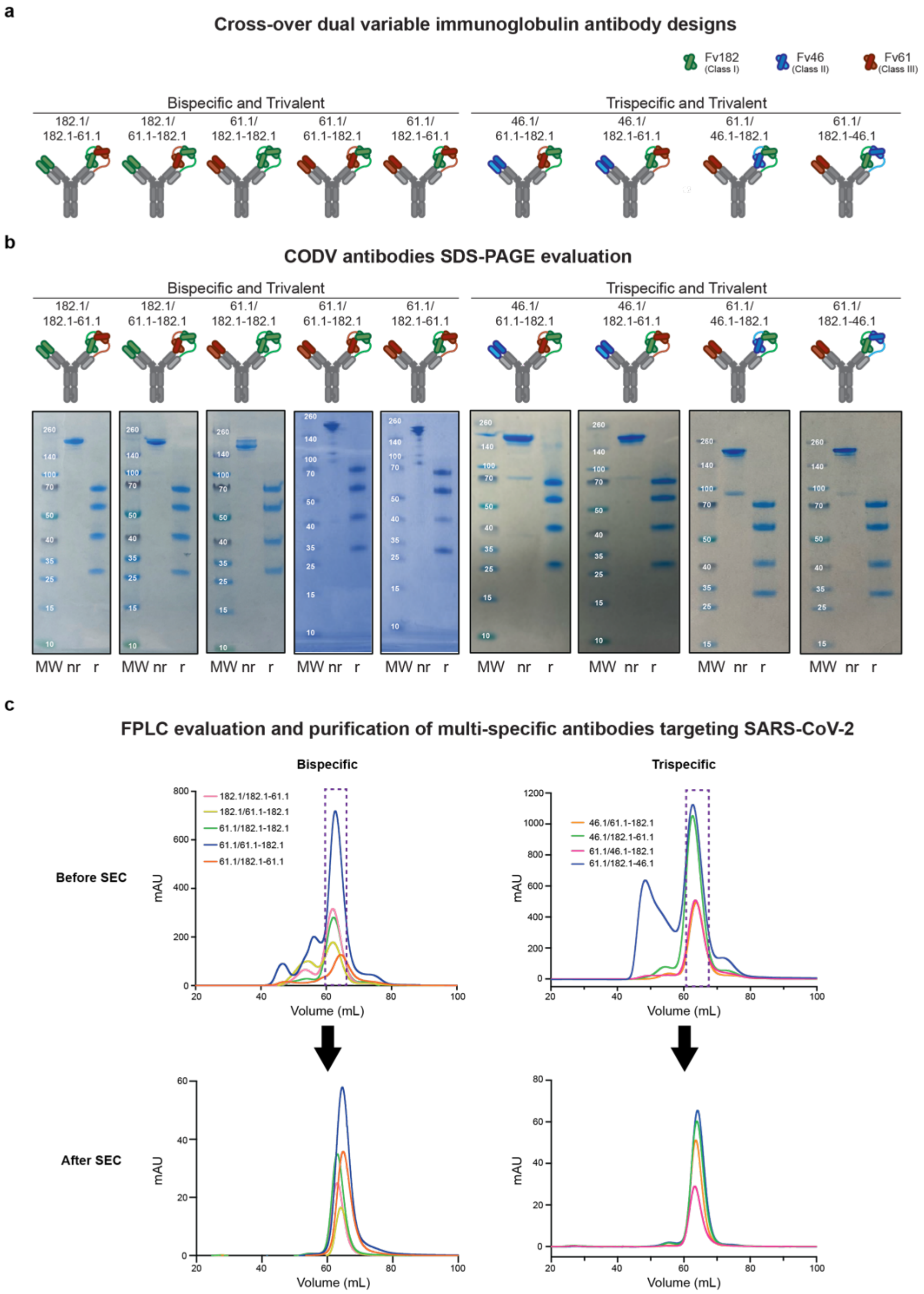


**Extended Data Figure 1. Production and purification of SARS-CoV-2 CODV antibodies**

**a,** Cross-over dual variable (CODV) immunoglobulin antibody designs utilizing variable fragments (Fv) from B1-182.1 (Fv182.1, green), A19-46.1 (Fv46.1, blue) and A19-61.1 (Fv61.1, red). Fc regions contain "knob and hole" feature to increase yield and correct association of heavy chains. Five bispecific trivalent molecules (182.1/182.1-61.1, 182.1/61.1-182.1, 61.1/182.1-182.1, 61.1/61.1-182.1, 61.1/182.1-61.1) and four trispecific trivalent molecules (46.1/61.1-182.1, 46.1/182.1-61.1, 61.1/46.1-182.1, 61.1/182.1-46.1) were designed. **b,** Purity of cross-over dual variable (CODV) immunoglobulin antibodies were evaluated under non-reducing (nr) and reducing (r) conditions on a Coomassie SDS-PAGE gel (representative gels shown). **c,** CODV immunoglobulin bispecific and trispecific antibody traces shown before (top row) and after size exclusion chromatography (SEC) (bottom row). The purple box indicates that fractions combined to make final preparations of the indicated multispecific antibodies. The bottom row shows analytic SEC traces for the purified multispecifics. Shown are representative traces from production runs of antibodies. Antibody properties and yields are shown in Extended Data Table 1.

| ID | Molecular Weight (kDa) | Isoelectric Point | Extinction Coefficient. ($M^{-1}$ $cm^{-1}$) | Yield (mg/L) |
|---|---|---|---|---|
| **182.1/182.1-182.1** | 173.1 | 8.38 | 267120 | 14.5 |
| **182.1/182.1-61.1** | 173.4 | 8.39 | 271465 | 3.94 |
| **182.1/61.1-182.1** | 173.4 | 8.39 | 271465 | 2.6 |
| **61.1/182.1-182.1** | 173.4 | 8.39 | 271465 | 4.37 |
| **61.1/61.1-182.1** | 173.7 | 7.81 | 275810 | 1.6 |
| **61.1/182.1-61.1** | 173.7 | 7.81 | 275810 | 9.6 |
| **46.1/61.1-182.1** | 173.7 | 8.32 | 280280 | 9.24 |
| **46.1/182.1-61.1** | 173.7 | 8.32 | 280280 | 16.41 |
| **61.1/46.1-182.1** | 173.7 | 8.32 | 280280 | 7.94 |
| **61.1/182.1-46.1** | 173.7 | 8.32 | 280280 | 17.72 |

**Extended Data Table 1. Biochemical properties and production yields following SEC purification for each SARS-CoV-2 CODV antibodies**

**a**

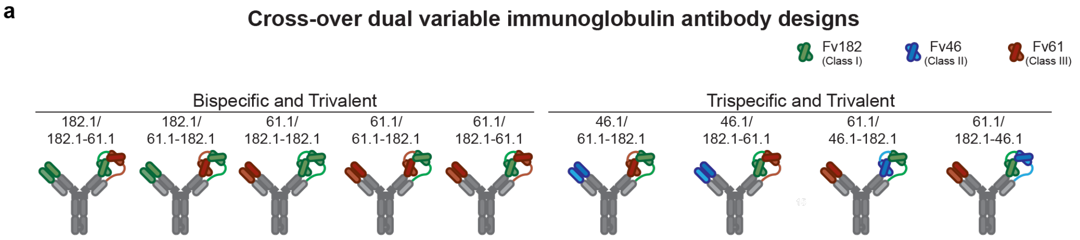


**b**

**Expected binding of Fv components to RBD mutations**

| | Fv182 | Fv46 | Fv61 |
|---|---|---|---|
| F486S | - | ++++ | ++++ |
| L452R | ++++ | - | ++++ |
| K444E | ++++ | ++++ | - |

**c**

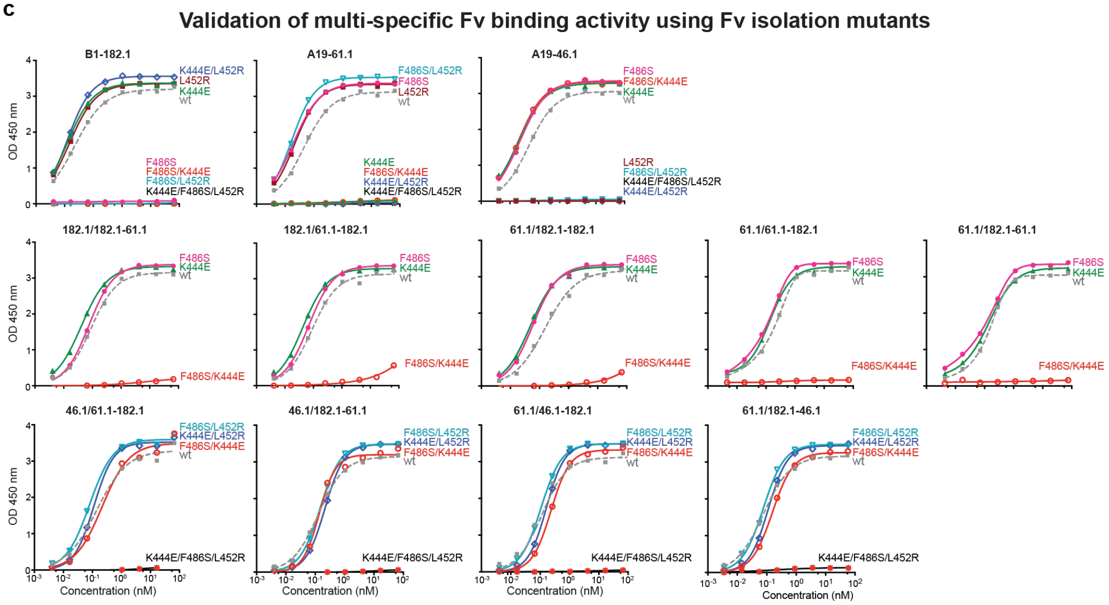


**d**

**Validation of multi-specific neutralization activity using Fv isolation virus mutants**

| | | D614G | | B1-182.1 Escape F486S/D614G | | A19-46.1 Escape L452R/D614G | | A19-61.1 Escape K444E/D614G | |
|---|---|---|---|---|---|---|---|---|---|
| | | $IC_{50}$ | $IC_{80}$ | $IC_{50}$ | $IC_{80}$ | $IC_{50}$ | $IC_{80}$ | $IC_{50}$ | $IC_{80}$ |
| | B1-182.1 | 15.7 | 34.9 | >67,000 | >67,000 | 25.0 | 75.8 | 11.1 | 29.9 |
| | A19-61.1 | 44.1 | 96.5 | 75.2 | 166 | 207 | 444 | >67,000 | >67,000 |
| | A19-46.1 | 113 | 339 | 133 | 390 | >67,000 | >67,000 | 472 | 1504 |
| Bispecific and Trivalent | 182.1/182.1-61.1 | 11.4 | 36.5 | 404 | 1320 | 24.1 | 113 | 7.0 | 56.1 |
| | 182.1/61.1-182.1 | 15.8 | 46.8 | 354 | 790 | 38.5 | 65.2 | 10.1 | 35.8 |
| | 61.1/182.1-182.1 | 67.4 | 128 | 320 | 1008 | 146 | 312 | 261 | 1090 |
| | 61.1/61.1-182.1 | 81.2 | 108.2 | 36.7 | 76.0 | 174.6 | 470.3 | >67,000 | >67,000 |
| | 61.1/182.1-61.1 | 20.2 | 46.4 | 29.4 | 59.0 | 93.8 | 230.2 | 3053.4 | 8211.7 |
| Trispecific and Trivalent | 46.1/61.1-182.1 | 48.7 | 119 | 112 | 243 | 201 | 603 | 189 | 487 |
| | 46.1/182.1-61.1 | 31.7 | 62.9 | 80.0 | 271 | 279 | 910 | 46.9 | 140 |
| | 61.1/46.1-182.1 | 30.8 | 48.4 | 68.6 | 164 | 233 | 934 | 42.0 | 140 |
| | 61.1/182.1-46.1 | 82.0 | 183 | 82.6 | 212 | 1010 | 2097 | 89.3 | 356 |

| pM |
|---|
| <10 |
| 10-100 |
| 100-1000 |
| 1000-10,000 |
| 10,000-67,000 |
| >67,000 |

**Extended Data Figure 2. Functional assessment and effect of position of Fv components against RBD knockout mutations**

**a,** Cross-over dual variable (CODV) immunoglobulin antibody designs utilizing variable fragments (Fv) from B1-182.1 (Fv182.1, green), A19-46.1 (Fv46.1, blue) and A19-61.1 (Fv61.1, red). Fc regions contain “knob and hole” feature to increase yield and correct association of heavy chains. Five bispecific trivalent molecules (182.1/182.1-61.1, 182.1/61.1-182.1, 61.1/182.1-182.1, 61.1/61.1-182.1, 61.1/182.1-61.1) and four trispecific trivalent molecules (46.1/61.1-182.1, 46.1/182.1-61.1, 61.1/46.1-182.1, 61.1/182.1-46.1) were designed. **b,** Shown are the expected binding of Fv182.1, Fv46.1 and Fv61.1 to RBD proteins containing the indicated Fv isolating mutations, that were selected to knockout binding a single component Fv while leaving binding of the remain Fv components unchanged. **c,** Validation of multispecific Fv binding activity using Fv isolating mutations in RBD and ELISA. Shown is ELISA binding for parental component antibodies controls (top row, B1-182.1, A19-46.1, A19-61.1), bispecific antibodies (middle row, 182.1/182.1-61.1, 182.1/61.1-182.1, 61.1/182.1-182.1, 61.1/61.1-182.1, 61.1/182.1-61.1) and trispecific antibodies (bottom row, 46.1/61.1-182.1, 46.1/182.1-61.1, 61.1/46.1-182.1 and 61.1/182.1-46.1) against WA-1 RBD (wt) or RBD proteins containing one or more Fv-specific mutations shown in panel B chosen to interrogate binding of one Fv component at a time in each multispecific. For example, F486S/K444E would show the binding of Fv46 component of trispecific antibodies because it does not allow binding of Fv182. and Fv61. **d,** Neutralization of candidate multispecific antibodies against D614G or mutants that target Fv182 (F486S/D614G), Fv46 (L452R/D614G) and Fv61 (K444E/D614G). Neutralization in pM is shown. Ranges are indicated with light blue (>67,000 pM), yellow (>10,000 to ≤67,000 pM), orange (>1000 to ≤10000 pM), red (>100 to ≤1000 pM), maroon (>10 to ≤100 pM), and purple (≤10 pM).

**Neutralization of B.1.1.7, BA.1, BA.2 and BA.4/5 by 46.1-182.1v/61.1-182.1v**

| | B.1.1.7 | | BA.1 | | BA.2 | | BA.4/5 | |
|---|---|---|---|---|---|---|---|---|
| | $IC_{50}$ | $IC_{80}$ | IC50 | IC80 | IC50 | IC80 | $IC_{50}$ | $IC_{80}$ |
| LY-CoV555 | 14.5 | 36.1 | >67,000 | >67,000 | >67,000 | >67,000 | >67,000 | >67,000 |
| 46.1-182.1v/61.1-182.1v | 18.7 | 71.0 | 28.5 | 87.6 | 9.7 | 25.0 | 1839 | 4366 |

| pM |
|---|
| <10 |
| 10-100 |
| 100-1000 |
| 1,000-10,000 |
| 10,000-67,000 |
| >67,000 |

**Extended Data Figure 3**. **46.1-182.1v/61.1-182.1v is a potently neutralizing tetravalent trispecific antibody**.

Neutralization of B.1.1.7, BA.1, BA.2 and BA.4/5 by 46.1-182.1v/61.1-182.1v. Neutralization in pM is shown. Ranges are indicated with light blue (>67,000 pM), yellow (>10,000 to ≤67,000 pM), orange (>1000 to ≤10000 pM), red (>100 to ≤1000 pM), maroon (>10 to ≤100 pM), and purple (≤10 pM).
